# MatchMaps: Non-isomorphous difference maps for X-ray crystallography

**DOI:** 10.1101/2023.09.01.555333

**Authors:** Dennis E. Brookner, Doeke R. Hekstra

## Abstract

Conformational change mediates the biological functions of macromolecules. Crystallographic measurements can map these changes with extraordinary sensitivity as a function of mutations, ligands, and time. The isomorphous difference map remains the gold standard for detecting structural differences between datasets. Isomorphous difference maps combine the phases of a chosen reference state with the observed changes in structure factor amplitudes to yield a map of changes in electron density. Such maps are much more sensitive to conformational change than structure refinement is, and are unbiased in the sense that observed differences do not depend on refinement of the perturbed state. However, even minute changes in unit cell properties can render isomorphous difference maps useless. This is unnecessary. Here we describe a generalized procedure for calculating observed difference maps that retains the high sensitivity to conformational change and avoids structure refinement of the perturbed state. We have implemented this procedure in an open-source python package, MatchMaps, that can be run in any software environment supporting PHENIX and CCP4. Through examples, we show that MatchMaps “rescues” observed difference electron density maps for poorly-isomorphous crystals, corrects artifacts in nominally isomorphous difference maps, and extends to detecting differences across copies within the asymmetric unit, or across altogether different crystal forms.

**Synopsis:** MatchMaps is a generalization of the isomorphous difference map allowing for computation of difference maps between poorly-isomorphous and non-isomorphous pairs of crystallographic datasets. MatchMaps is implemented as a simple-to-use, python-based command-line interface.

## 1. Introduction

X-ray crystallography provides a powerful method for characterizing the changes in protein structure caused by a perturbation (Hekstra *et al*., 2016; Keedy *et al*., 2018; Bhabha *et al*., 2015; Brändén & Neutze, 2021). For significant structural changes, it is usually sufficient to refine separate structural models for each dataset and draw comparisons between the refined structures. However, for many conformational changes, coordinate-based comparisons are inaccurate and insensitive.

In crystallography, electron density is not observed directly, but rather a diffraction pattern consisting of reflections with intensities proportional to the squared amplitudes of the structure factors—the Fourier components of the electron density. Unfortunately, the phases of these structure factors are not observable. These phases correspond in real space to shifts of the sinusoidal waves that add up to an electron density pattern. Accordingly, phases are usually calculated from a refined model. Since phases have a strong effect on the map appearance (Read, 1986), naive electron density maps calculated using observed amplitudes and model-based phases will tend to resemble the model, a phenomenon known as model bias.

The gold standard for detecting conformational change in crystallographic data is the isomorphous difference map(Rould & Carter, 2003). An isomorphous difference map is computed by combining differences in observed structure factor amplitudes with a single set of phases. The phases are usually derived from a model for one of the two states, chosen as a reference. Thus, difference density Δ*ρ*(**x**) is approximated as:.

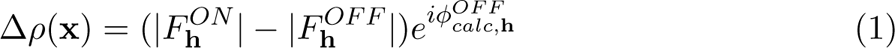

where |*F*_h_^*ON*^| and |*F*_h_^*OFF*^| are sets of observed structure factor amplitudes from the ON (perturbed) and OFF (reference) datasets respectively, 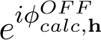 is a set of calculated structure factor phases derived from a structural model of the OFF data, **h** is shorthand for the triplet of Miller indices (*h, k, l*), and **x** is shorthand for the realspace point (*x, y, z*). Crucially, therefore, isomorphous difference maps do not include any information derived from modeling of the ON structure. Any difference electron density relating to the ON data relative to the OFF data (e.g., positive difference density for a bound ligand) is thus guaranteed not to be biased by previous modeling of the ON state. Unfortunately, interesting conformational changes often slightly alter the packing of molecules in the unit cell, which can manifest as changes in unit cell dimensions. Unit cells constants are also sensitive to temperature(Fraser *et al*., 2011), radiation damage(Ravelli & McSweeney, 2000), pressure(Barstow *et al*., 2008), and humidity(Farley *et al*., 2014), meaning that even data collected on the same crystal may not be quite isomorphous.

In this contribution, we will illustrate the consequences of deviations from perfect isomorphism, introduce an approach to the calculation of difference maps without perfect isomorphism, and describe examples of the application of the software implementing this approach (MatchMaps) to a number of typical use cases. We find the MatchMaps approach, moreover, to be applicable to molecules related by noncrystallographic symmetry and molecules crystallized in altogether different crystal forms.

### 1.1. Implications of isomorphism

We begin by demonstrating the consequences of small deviations from perfect isomorphism. Our example makes use of three datasets, all of *E. coli* dihydrofolate reductase (DHFR) crystallized in spacegroup *P* 2_1_2_1_2_1_. These datasets vary by which ligands are bound to DHFR; we will discuss these ligands further below. Datasets 1RX2 and 1RX1 have unit cell dimensions identical to within 0.4%, whereas datasets 1RX2 and 1RX4 differ by 2% along the *c*-axis. Reflections in diffraction experiments report on different 3D frequency components of the electron density of molecules in unit cell. As such, the shape of the molecular arrangement may look essentially the same (that is, isomorphous) at low spatial resolution, yet entirely different at high resolution (recall that the contributions of different atoms, *j*, to structure factors add up by terms exp (2*π***s** · **r***_j_*), for scattering vector **s** and atomic position **r***_j_*). Measuring this quantitatively, we see much higher correlations between the structure factor amplitudes for our highly isomorphous pair of datasets (1RX2 and 1RX1) than for our “poorly-isomorphous” pair (1RX2 and 1RX4) (solid lines in Figure 1a). We find a similarly stark difference in correlations for the phases of refined models, whether measured by a figure of merit, ⟨cos (*ϕ*_2_ − *ϕ*_1_)⟩, or by a correlation coefficient (liable to small phase wrapping artifacts). The loss in similarity of phases is visually striking (Figure 1, panels b and c).

**Figure 1.**
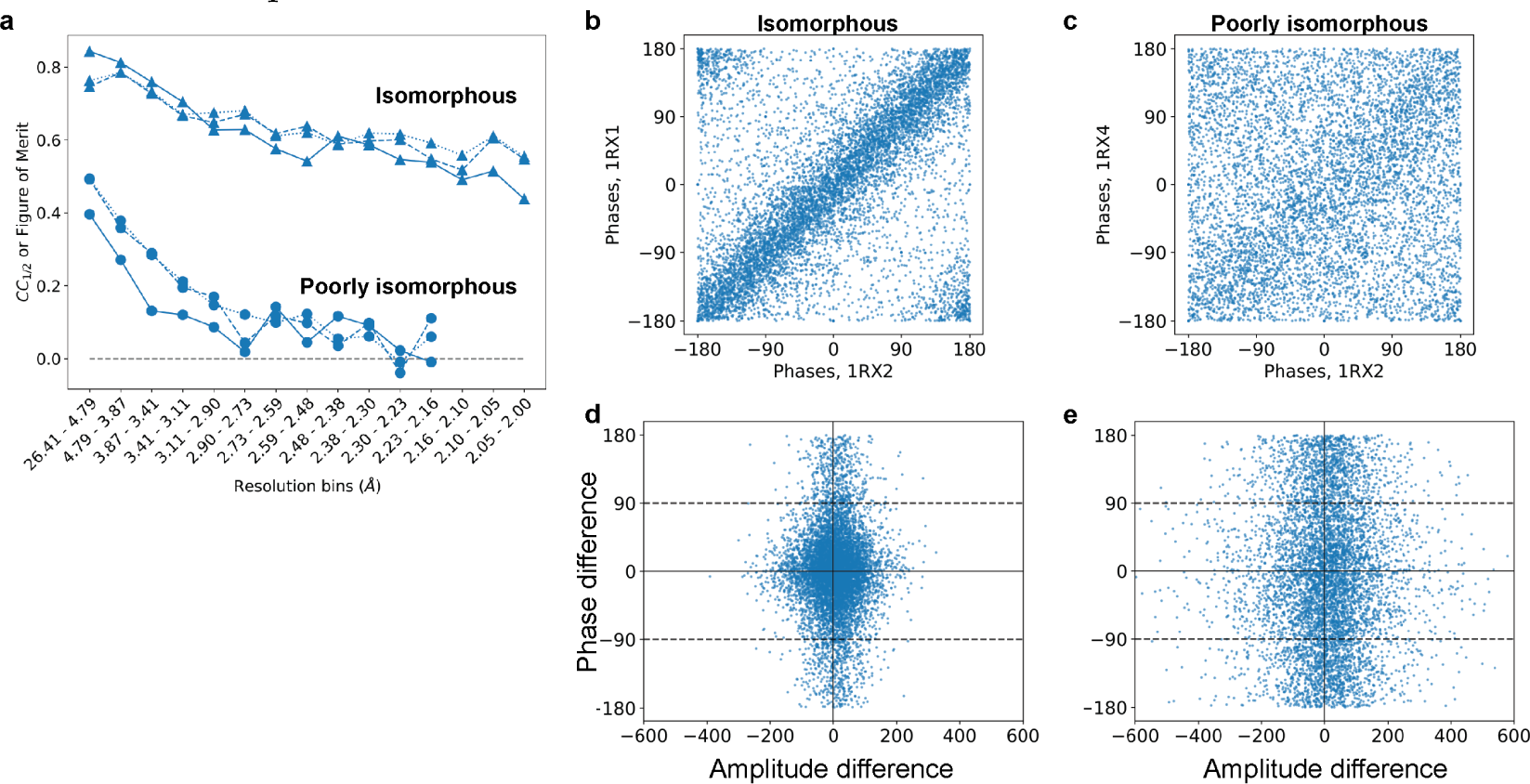
Structure factors depend sensitively on isomorphism. Here, we compare the *E. coli* DHFR dataset 1RX2 with a highly isomorphous structure (1RX1) and with a poorly isomorphous structure (1RX4, see also figure 2). (a) Correlation coefficients for the isomorphous pair (circles) and poorly isomorphous pair (triangles): correlations were computed between structure factor amplitudes (solid lines) and cosines of structure factor phases (dashed lines), aggregated per resolution bin. Additionally, a figure of merit (mean of cosine of differences) was computed between corresponding phases (dotted lines). While the isomorphous data correlate well even at high resolution, the poorly isomorphous data are uncorrelated even at moderate resolution. (b, c) Structure factor phases appear (b) highly correlated for the isomorphous structures, but (c) mostly uncorrelated between poorly-isomorphous structures. (d, e) For an isomorphous difference map to be meaningful, structure factor amplitudes should only differ when the phase difference is small, and structure factor phases should only differ when the amplitude difference is small (Rould & Carter, 2003). This requirement is met in the isomorphous case (d), but not in the non-isomorphous case (e). All panels: Computed phases are obtained from the “PHIC” column of the deposited MTZ files.

We expect the consequences of such loss of isomorphism to be severe: the computation of an isomorphous difference map requires that 1) amplitude differences are large only when phase differences are small, and conversely that 2) phase differences are large only when amplitude differences are small. These requirements follow from Equation 1 above, and are depicted visually in (Rould & Carter, 2003). The isomorphous data meet these requirements (Figure 1d). In contrast, the poorly-isomorphous datasets display consistently large structure factor amplitude differences, regardless of the corresponding structure factor phase difference (Figure 1e).

### 1.2. Rethinking isomorphous difference maps via the linearity of the Fourier transform

An isomorphous difference map is typically computed by first subtracting the structure factor amplitudes (e.g., subtracting in reciprocal space) and then applying the Fourier transform to convert the structure factor differences into a real-space difference map. However, because the Fourier transform and subtraction are both linear operations, their order can be switched without changing the result: one might just as well calculate two electron density maps first and then subtract those maps voxel-by-voxel to yield an isomorphous difference map.

This reordering suggests how difference map computation can be generalized beyond the isomorphous case. Specifically, we see that the step in the algorithm most specific to the assumption of isomorphism is the construction of “hybrid” structure factors, which combine the observed structure factor amplitudes for the ON data |(*F*_*obs,h*_^*ON*^)| with the calculated structure factor phases for the OFF data (*ϕ_calc,h_^OFF^*). The resulting structure factors thus have the form:

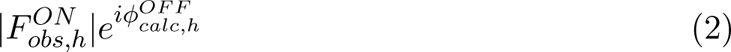

Critically, if the ON and OFF data differ in unit cell volume and/or molecular orientation, these OFF phases may be incompatible with the ON amplitudes.

The method presented below improves these “hybrid” structure factors by computing phases that account for the (generally uninteresting) shifts in molecular position and orientation without removing any signal associated with “interesting” changes.

## 2. The MatchMaps algorithm

The goal of MatchMaps is to achieve the best possible real-space difference density map without utilizing a prior model of any structural changes of interest. To compute a real-space difference density map, one first needs to approximate structure factor phases for each dataset. As discussed above, the isomorphous difference map makes the simplifying assumption that the same set of structure factor phases can be used “as is” for both structures.

The key to MatchMaps is to improve phases for the ON data via rigid-body refinement of the OFF starting model against the ON structure factor amplitudes. This rigid-body refinement step improves phases by optimally placing the protein model in space. However, the restriction of this refinement to only whole-model rigid-body motion protects these new phases from bias towards modeled structural changes. The result is two sets of complex structure factors which make use of the information encoded in the structure factor amplitudes without relying on a second input model. Next, each set of complex structure factors is Fourier-transformed into a real-space electron density map. These two real-space maps will not necessarily overlay in space. However, the rotation and translation necessary to overlay the maps can be obtained from the results of rigid-body refinement. Following real-space alignment, the maps can be subtracted voxel-wise to compute a difference map.

In the idealized case—similar structures, oriented identically in space, with identical unit cells—MatchMaps will perform essentially identically to an isomorphous difference map. However, as we show in the examples below, MatchMaps is more capable than a traditional isomorphous difference map of handling datasets that diverge from this ideal. Furthermore, even in seemingly simple cases where isomorphous difference maps perform well, the real-space MatchMaps approach can show distinct improvements.

### 2.1. Details of algorithmic implementation

The full MatchMaps algorithm is as follows. As inputs, the algorithm requires two sets of structure factor amplitudes (referred to as ON and OFF datasets, for simplicity) and a single starting model (corresponding to the OFF data).

1. If necessary, place both sets of structure factor amplitudes on a common scale using CCP4 (Agirre *et al*., 2023)’s `SCALEIT` (Henderson & Moffat, 1971) utility.
2. Truncate both datasets to the same resolution range. This prevents the final difference map from preferentially displaying high-resolution features from the higher-resolution dataset.
3. Generate phases for each dataset via the `phenix.refine` (Liebschner *et al*., 2019) program. For each dataset, the OFF starting model is used, and only rigid-body refinement is permitted to prevent the introduction of model bias. Bulk-solvent scaling may be either included (by default) or omitted from refinement. Including bulk-solvent scaling leads to better refinement statistics and higher map quality overall. However, bulk-solvent scaling may “flatten” desired signal in the solvent region, e.g. for a large bound ligand. This trade-off is left to the user.
4. Create complex structure factors by combining observed structure factor amplitudes with computed structure factor phases obtained from refinement. Fouriertransform each set of complex structure factors into a real-space electron density map; this is performed using the python packages `reciprocalspaceship` (Greisman *et al*., 2021) and `gemmi` (Wojdyr, 2022).
5. Compute the translation and rotation necessary to overlay the two rigid-body refined models. Apply this translation-rotation to the ON real-space map such that it overlays with the OFF map. These computations are carried out using `gemmi`. Note that the two rigid-body refined models are identical aside from translation and rotation, rendering trivial the atom selection for alignment.
6. Subtract real-space maps voxel-wise.
7. Apply a solvent mask to the final difference map.

We note that MatchMaps is structured such that step 2 can be generalized to not only rigid-body refinement but refinement of any “uninteresting features”, if the user provides a custom PHENIX parameter file as specified in the online documentation. For example, if the starting model contains multiple protein chains, each chain can be rigid-body-refined separately.

### 2.2. Installation

MatchMaps can be installed using the `pip` python package manager (`pip install matchmaps`). The various pure-python dependencies of MatchMaps are handled by `pip`. Additionally, MatchMaps requires installation of the popular CCP4 and Phenix software suites for crystallography. Once installed, the above protocol can be run in a single step from the command line.

In addition to the base MatchMaps command-line utility, the utilities `matchmaps.ncs` and `matchmaps.mr` provide additional functionalities explored in the examples below and the online documentation. MatchMaps is fully open-source and readily extensible for novel use cases.

For more information, read the MatchMaps documentation at: https://rs-station.github.io/matchmaps.

## 3. Comparison of MatchMaps with alternative approaches

Of course, MatchMaps is not the only possible method for comparing subtle structural changes as differences in electron density. Two possible alternate methods contrast interestingly with MatchMaps, and these methods warrant discussion here.

### 3.1. F_o_ − F_c_ difference maps

A common element of structure refinement is the so-called “*F_o_* − *F_c_*” map (or, often, *mF_o_*−*DF_c_*), which is used to describe how the modeled structure differs from the data. Details of the construction of such a map can be found elsewhere (Lamb *et al*., 2015). In practice, *F_o_* − *F_c_*maps are often the output of a procedure including refinement of atomic coordinates. In principle, however, an *F_o_* − *F_c_*map can derive from a rigid-body-only refinement of a known structure to a new dataset. In this latter scenario, the *F_o_* − *F_c_* map is similar to a MatchMaps difference map (or in an isomorphous case, an isomorphous difference map).

The difference between an *F_o_* − *F_c_*map and a MatchMaps difference map is that whereas MatchMaps only ever uses *observed* structure factor amplitudes, the *F_o_* − *F_c_*map describes the OFF/reference dataset using *calculated* structure factor amplitudes. In the limiting case where the OFF model describes the OFF data perfectly, the *F_o_*− *F_c_*map should look like a MatchMaps difference map. In fact, an *F_o_* − *F_c_*map may look better, because the map coefficients include only one set of measurement errors. Unfortunately, however, any modeling errors of the OFF/reference state will be included the final difference map. Accordingly, in an *F_o_* − *F_c_*map, it is impossible to distinguish “real signal” (differences between the ON and OFF data) from modeling errors. We illustrate this undesired behavior below.

Note that the map coefficients for an *F_o_* − *F_c_* map are created and saved by MatchMaps (if the `--keep-temp-files` flag is used), facilitating easy comparison between these two map types if desired.

### 3.2. PanDDA

A recent, popular method for extracting subtle ligand-binding signal from crystallographic data is the Pan-Dataset Density Analysis (PanDDA) approach (Pearce *et al*., 2017). A key practical difference between PanDDA and MatchMaps is that while PanDDA expects several (typically ∼dozens) datasets, MatchMaps supports only two datasets at once. Additionally, whereas MatchMaps never changes internal atomic coordinates of the input model, PanDDA aligns all input structures and maps via a local warping procedure. Thus, PanDDA reduces its ability to describe protein conformational changes in order to maximize its ability to detect weak ligand-binding events.

## 4. Examples

The following examples explore the benefits and functionalities offered by MatchMaps. All examples make use of published crystallographic data available from the Protein Data Bank. Scripts and data files for reproducing the figures can be found on Zenodo.

### 4.1. MatchMaps for poorly-isomorphous datasets

The enzyme Dihydrofolate Reductase (DHFR) is a central model system for understanding the role of conformational change in productive catalytic turnover(Sawaya & Kraut, 1997; Boehr *et al*., 2006; Bhabha *et al*., 2011). Specifically, the active-site Met20 loop of *E. coli* DHFR can adopt several different conformations, each stabilized by specific bound ligands and crystal contacts(Sawaya & Kraut, 1997). DHFR bound to NADP^+^ and substrate analog folate adopts a “closed” Met20 loop (PDB ID 1RX2), whereas DHFR bound to NADP^+^ and product analog (dideazatetrahy-drofolate) adopts an “occluded” Met20 loop (PDB ID 1RX4). These structures are highly similar, other than the relevant changes at the active site (Figure 2a, structural changes shown in boxes; RMSD 0.37 Å for protein C-alpha atoms excluding Met20 loop).

**Figure 2:**
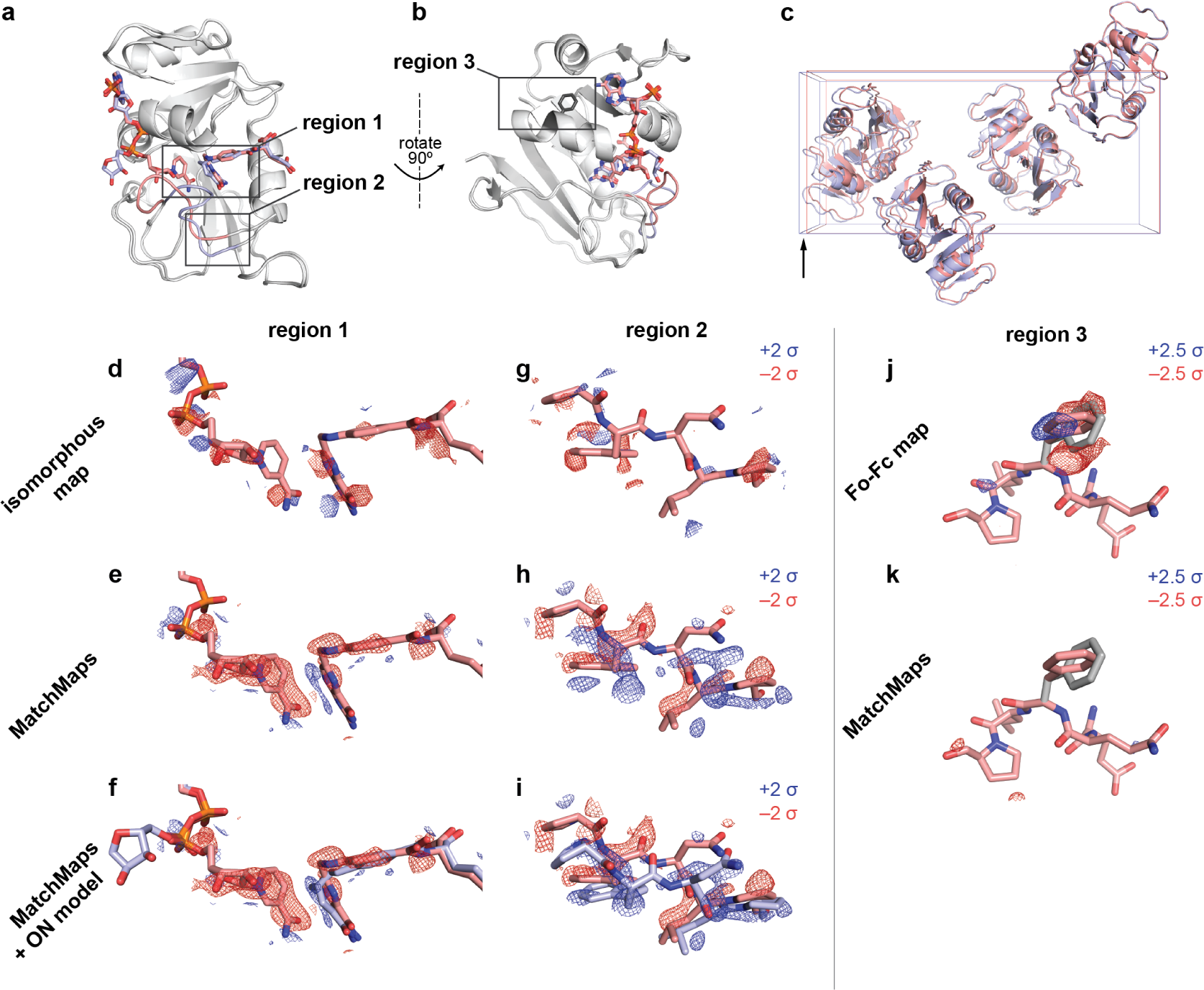
MatchMaps for poorly-isomorphous structures. (a) The structures 1RX2 and 1RX4 are similar overall (gray cartoons). The structures differ mainly at the active site loop (1RX2, pink cartoon; 1RX4, blue cartoon) and in the positions of the active-site ligands (1RX2, pink sticks; 1RX4, blue sticks). (b) Same as (a), but rotated 90 degrees to the right about the vertical axis (pictured). Sidechain for phenylalanine 103 is shown as dark gray sticks. (c) The unit cells of 1RX2 and 1RX4 differ by 2% along the longest dimension (left to right in this figure) from 98.912 Åto 100.879 Å. (d-f) Visualization of the change in ligand position between 1RX2 (red sticks) and 1RX4 (f, blue sticks). Positive difference density is shown as blue mesh; negative different density is shown as red mesh. Importantly, the 1RX4 structural coordinates were not used in the creation of the the isomorphous or MatchMaps maps. The isomorphous difference map contains essentially no interpretable signal. In contrast, the MatchMaps map (e-f) contains clear signal for disappearance of the cofactor and the lateral sliding of the substrate. (f) is the same as (e), with the addition of the 1RX4 structural coordinates as blue sticks. (g-i) Visualization of the change in loop conformation between 1RX2 and 1RX4. Only protein residues 21-25 are shown. Coloring is as in panels d-f. (g) Again, the isomorphous difference map is not interpretable. (h-i) The MatchMaps positive difference density clearly corresponds with the 1RX4 structural model, which was not used in the creation of the map. (i) is the same as (h), with the addition of the 1RX4 structural coordinates as blue sticks. (j-k) Impact of a spurious conformer on *F_o_* − *F_c_*, MatchMaps. 1RX2 model for residues 101-105 are shown as pink sticks. The spurious conformer for Phe103 is shown as gray sticks. (j) *F_o_* − *F_c_* map shows clear positive (blue) and negative (red) density recognizing the erroneous conformer as a conformational change. (k) MatchMaps does not show difference density for the spurious conformer.

Importantly, the presence of the occluded loop conformation leads to altered crystal packing wherein the crystallographic *b* axis increases by 2%, from 98.91 Å to 100.88Å (Figure 2c). Thus, 1RX2 and 1RX4 are “poorly isomorphous”; this means that these structures, though extremely similar, cannot be effectively compared by an isomorphous difference map (Figure 2d,g). We illustrated the striking change in phases between these structures in Figure 1. MatchMaps is able to account for this poor isomorphism and recover the expected difference signal.

First, we focused on ligand rearrangement in the active site. In the occluded-loop structure, the cofactor (Figure 2d-f, left) leaves the active site while the substrate (Figure 2d-f, right) slides laterally within the active site. MatchMaps shows this expected signal, with negative (red) difference density for the cofactor and paired positive (blue) and negative (red) difference density for the substrate (Figure 2e-f). By contrast, an isomorphous difference map (Figure 2d) is unable to recover this signal. A model of the occluded-loop structure is shown for clarity in Figure 2f as blue sticks and clearly matches with the positive difference density. Importantly, this ON model is never used in the computation of the MatchMaps map.

We find a similar result around residues 21-25 of the Met20 loop (Figure 2g-i). Again, MatchMaps shows readily interpretable difference signal for the change in loop conformation between the closed-loop (red) and occluded-loop (blue) structures (Figure 2h-i). The isomorphous difference map, on the other hand, contains no interpretable signal in this region of strong structural change (Figure 2g). The occluded-loop model is shown for visual comparison in Figure 2i but was not used for computation of the MatchMaps map.

#### 4.1.1. MatchMaps is not susceptible to modeling errors

As discussed above, *F_o_*− *F_c_*maps can often display similar information to MatchMaps difference maps. Indeed, an *F_o_* − *F_c_*map can perform similarly to MatchMaps in this case (Figure S1). However, *F_o_* − *F_c_* maps will inevitably also contain signal that is not in fact a difference between ON and OFF datasets, but rather is a modeling error of the OFF model to the OFF data. We demonstrate this behavior by introducing a spurious conformer of phenylalanine 103 to the OFF starting model used above. Phe103 lies in a region distal to the ligands and active site (Figure 2b). An *F_o_* − *F_c_*map, which inherently includes modeling errors, shows strong positive and negative difference density suggesting the correct Phe103 conformer (Figure 2j). From the *F_o_* − *F_c_* alone, it would be impossible to determine if this signal represented a difference between the ON and OFF data or a modeling error. In contrast, the MatchMaps difference map shows no difference density for this sidechain (Figure 2k). This is the desired and expected result; neither dataset’s *F_o_*contains any information about this spurious conformer.

### 4.2. MatchMaps corrects artifacts even for isomorphous datasets

The enzyme Protein Tyrosine Phosphatase 1B (PTP1B) plays a key role in insulin signaling(Elchebly *et al*., 1999), making it a long-standing target for the treatment of diabetes using ortho- and allosteric drugs (Wiesmann *et al*., 2004; Keedy *et al*., 2018; Choy *et al*., 2017). For illustration, we compare recent high-quality room-temperature structures of the apo protein (PDB ID 7RIN) with the protein bound to the competitive inhibitor TCS401 (PDB ID 7MM1)(Greisman *et al*., 2022). In addition to the presence/absence of signal for the ligand itself, the apo structure exhibits an equilibrium between “open” and “closed” active-site loops(Whittier *et al*., 2013), whereas the bound structure shows only the closed loop.

The datasets 7RIN and 7MM1 are sufficiently isomorphous that an isomorphous difference map reveals the main structural changes. MatchMaps performs similarly. Strong positive difference density (blue mesh) is seen for the TCS401 ligand (grey sticks) in both the isomorphous difference map (Figure 3c) and the MatchMaps difference map (Figure 3d). Around residues 180-182 of the active-site loop (known as the WPD loop), both the isomorphous difference map (Figure 3e) and MatchMaps difference map (Figure 3f) show strong signal for the decrease in occupancy (red mesh) of the open loop conformation (red sticks) and an increase in occupancy (blue mesh) of the closed loop conformation (blue sticks).

**Figure 3:**
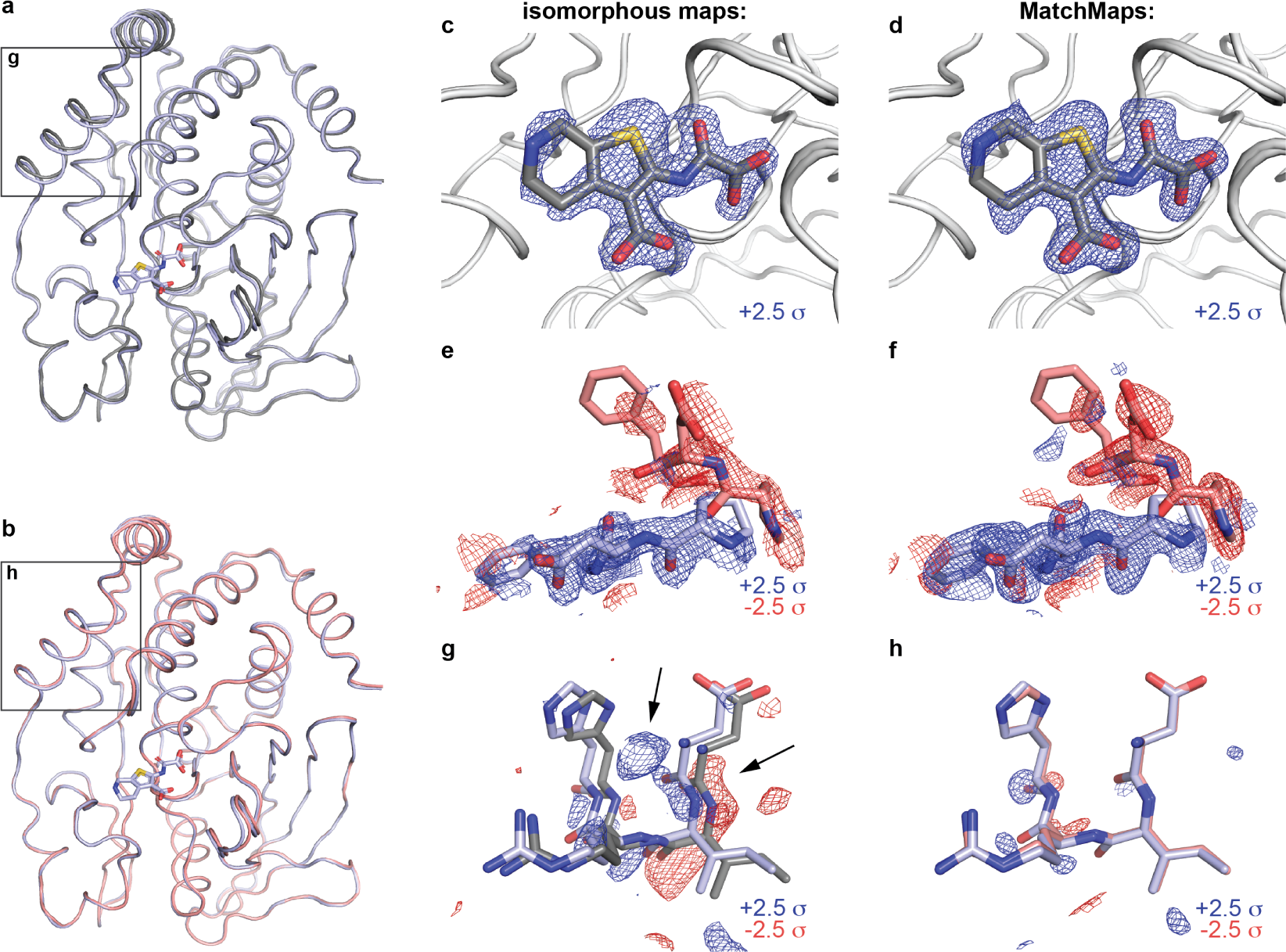
MatchMaps removes artifact even in isomorphous case. Comparison of apo (7RIN) and TCS401-bound (7MM1) structures of protein tyrosine phosphatase 1B. (a) The apo (gray) and bound (blue) structural models overlay well, but differ by a slight rotation. The difference in the models is especially apparent in the boxed region (see g). (b) Aligning the apo model (red) to the bound model (blue) reveals that the structures overlay even better than the original coordinates suggest (see h). (c-d) Both the isomorphous map (c) and MatchMaps map (d) are able to clearly show the bound ligand. The TCS401 ligand (gray sticks) is shown for clarity but was not included in the computation of either map. Positive difference density is shown as a blue mesh. (e-f) Closeup of residues 180-182. Similarly to (c-d), the change in loop equilibrium between open (red mesh, red sticks) and closed (blue mesh, blue sticks) is apparent in both maps. (g-h) Residues 22-25 are shown. Though these structures meet the requirements for isomorphism, the refined protein models still differ by a slight rotation. (g) The isomorphous difference map recognizes the artifactual difference between 7MM1 (blue) and 7RIN (gray) model locations, which manifests as strong difference signal. This artifact is comparable in magnitude to the “true” signal in panels (c) and (e). (h) MatchMaps internally aligns the models before subtraction and therefore avoids this artifact. The bound model after alignment to the apo model is shown in red. At +/- 2.5 σ, there is no significant signal in the MatchMaps map for this region.

However, even in this seemingly straightforward case, we find that the isomorphous difference map is susceptible to an artifact resulting from a slight (1.37 degrees) rotation of the protein. The displacement between the original refined structural coordinates of each structure is especially strong around residues 22-25 (Figure 3a, boxed region; Figure 3g, apo model in gray, bound model in blue). In this region, an isomorphous difference map picks up on this artifactual difference between the datasets and displays strong difference signal (blue and red mesh). Remarkably, this signal is similar in magnitude to the “true” signal seen in panels 3c and 3e. In contrast, MatchMaps internally aligns the models after the computation of phases and before subtraction. Figure 3b (boxed region) and Figure 3h (apo model in red, bound model in blue) show residues 22-25 following whole-molecule alignment of the protein models. Following global alignment of the refined models, it is clear that this region does not contain any “interesting” signal. Sure enough, the MatchMaps difference map contains no strong signal in this region. In fact, the faint signal that persists in the MatchMaps map for this region seems to report on a slight remaining coordinate displacement in this region following whole-molecule alignment.

### 4.3. matchmaps.mr: comparing data from different spacegroups

For many protein systems, careful analysis of electron density change is stymied for pairs of similar structures which crystallize in different crystal forms. The MatchMaps algorithm can be further generalized to allow comparison of datasets in entirely different crystal packings or spacegroups. Specifically, the OFF model can serve as a search model for molecular replacement for the ON data. Following this extra step, the algorithm proceeds identically. We implement this modified algorithm in the commandline utility `matchmaps.mr`.

One such example is the enzyme DHFR, which has been crystallized in many spacegroups(Sawaya & Kraut, 1997). Here, we examine two structures of the enzyme bound to NADP^+^, in spacegroups *P* 2_1_2_1_2_1_ (PDB ID 1RX1) and *C*2 (PDB ID 1RA1), visualized in Figure 4a. These structures are overall similar, but differ in the active site (Figure 4b-d). Here, we visualize these structural changes directly in electron density without introducing model bias.

**Figure 4:**
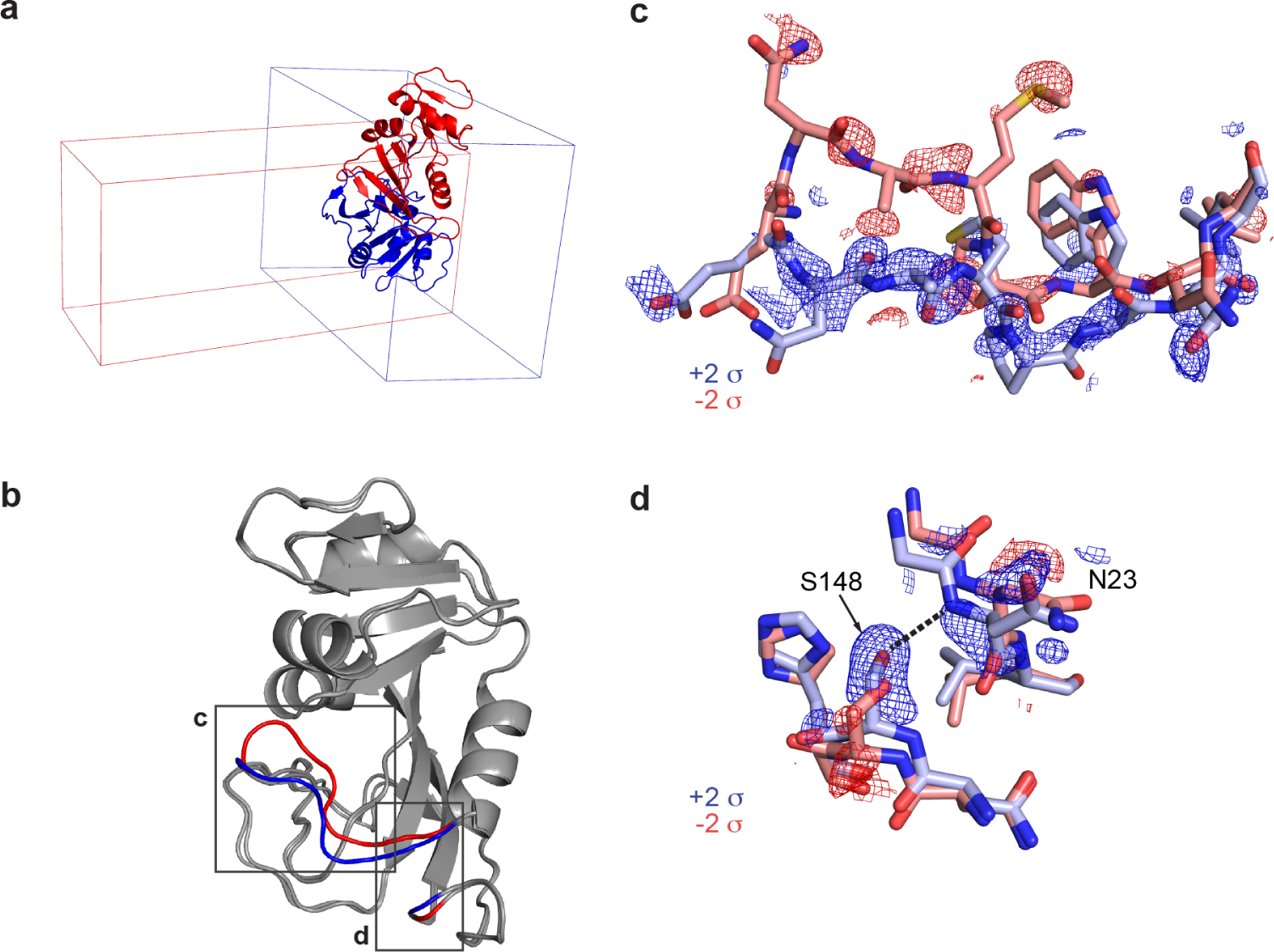
MatchMaps for difference maps across difference spacegroups. A variant of MatchMaps (implemented in the command line as matchmaps.mr) can be used to compute difference maps between two crystallographic datasets in entirely different spacegroups. (a) Overlay of structural models of DHFR in space-group *P* 2_1_2_1_2_1_ (PDB ID 1RX1, blue cartoon) and spacegroup *C*2 (PDB ID 1RA1, red cartoon) along with the respective unit cells for each. (b) Alignment of structures in *P* 2_1_2_1_2_1_ and *C*2 shows global agreement of structures. Structural differences are localized to the active site (boxed regions, *P* 2_1_2_1_2_1_ structure in red, *C*2 structure in blue) and are known to result from differences in crystal packing. (c) Closeup on residues 17-24. MatchMaps positive (blue) and negative (red) difference density clearly correspond to the refined structural coordinates for the *P* 2_1_2_1_2_1_ (red) and *C*2 (blue) models. Remarkably, the positive difference density is strong and clearly corresponds to the *C*2 structure, despite the *C*2 structure never being used in the creation of the map. (d) Closeup on the hydrogen bond between residues Ser148 and Asn23, which is only present in the *C*2 crystal form (blue sticks). MatchMaps (positive) difference density clearly indicates the hydrogen-bond-capable conformation.

Specifically, in the *P* 2_1_2_1_2_1_ structure, the active site Met20 loop adopts a closed conformation. In the *C*2 structure, the Met20 loop adopts an “open” conformation, which is stabilized by a crystal contact in this crystal form(Sawaya & Kraut, 1997). The difference between the open and closed loops is exemplified by residues 17-24 (Figure 4c). The open loop is stabilized by the formation of a key hydrogen bond between the Asn23 backbone and the Ser148 sidechain. In the closed conformation, Asn23 is too far from Ser148 to form a hydrogen bond (Figure 4d).

Remarkably, the positive difference density (blue) for the closed loop is strong and readily interpretable in Figures 4c-d. The MatchMaps map was computed only using the *P* 2_1_2_1_2_1_ (red) closed-loop model. This means that the signal for the open loop conformation is derived only from the observed structure factor amplitudes for the open-loop state in an unrelated crystal form!

### 4.4. matchmaps.ncs: comparing NCS-related molecules

The real-space portion of the MatchMaps algorithm can be repurposed to create “internal” difference maps across non-crystallographic symmetry (NCS) operations. As an example, we examined the crystal structure of the fifth PDZ domain (PDZ5) from the *Drosophila* protein Inactivation, no after-potential D (INAD). This domain plays an essential role in terminating the response of photoreceptors to absorbed photons by modulation of its ability to bind ligands (Mishra *et al*., 2007). In particular, the binding cleft of PDZ5 can be locked by formation of a disulfide bond between residues C606 and C645. PDZ5 was found to crystallize in a form with three molecules in the asymmetric unit (Figure 5a) where each molecules adopts a different state. Specifically, chain C contains a disulfide bond between residues Cys606 and Cys645, whereas chain B does not. Chains B and C overlay well other than the disulfide bond region (Figure 5b). Chain A adopts a bound state by binding the C terminus of chain C (not shown). MatchMaps enables calculation of an internal difference map, yielding a clearly interpretable difference map for the formation of the disulfide bond (Figure 5c).

**Figure 5:**
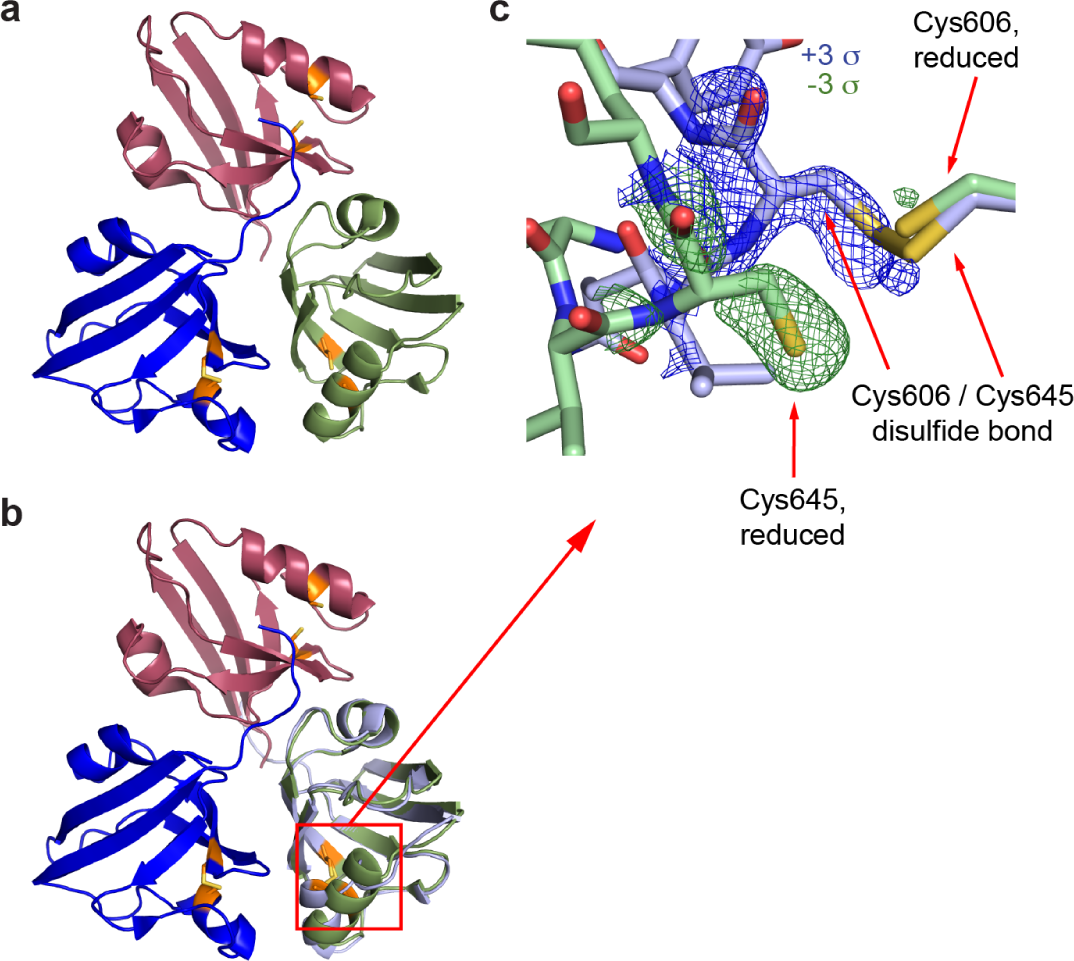
MatchMaps for internal difference maps across non-crystallographic symmetry operations. A variant of MatchMaps (implemented in the command line as `matchmaps.ncs`) can be used to compute internal difference maps across an NCS operation. (a) Overview of the three PDZ domains related by non-crystallographic symmetry. Chain A is shown in red, chain B in green, and chain C in blue. Residues Cys606 and Cys645, which can form a disulfide bond, are shown in orange. Coloring matches figure 2C from ref (Mishra *et al*., 2007). (b) Same as (a), plus a copy of chain C aligned and superimposed onto chain B, shown in light blue. (c) Zoom on the disulfide bond formation. Chain C (light blue sticks) contains a disulfide bond between Cys606 and Cys645, whereas chain B (green sticks) does not. The positive (blue) and negative (green) difference density corresponding to each chain is clearly visualized by MatchMaps.

## 5. Discussion

The isomorphous difference map has been the gold standard for detecting conformational change for many years (Henderson & Moffat, 1971; Rould & Carter, 2003). However, we show above that the same inputs—one structural model and two sets of structure factor amplitudes—can be combined to compute a difference map that shares the strength of an isomorphous difference map while ameliorating a key weakness. Specifically, structure factor phases are highly sensitive not only to structural changes (“interesting” signal) but also to changes in unit cell dimensions and model pose (“uninteresting” signal). The introduction of rigid-body refinement minimizes the contribution of this uninteresting signal to the final difference map. In Figure 2, we illustrate a case where a loss of isomorphism significantly degrades the signal of an isomorphous difference map. In this case, MatchMaps is still able to recover the expected difference signal. Figure 3 shows a situation where the isomorphous difference map performs well for the strongest difference signal. However, even in this seemingly straightforward use case, the isomorphous difference map is still susceptible to “uninteresting signal”. MatchMaps removes this artifact successfully.

In our experience, crystallographic perturbation experiments are often shelved due to changes in unit cell constants. MatchMaps removes, in principle, the requirement for isomorphism and allows for the analysis of far more crystallographic differences.

Furthermore, the computation of an isomorphous difference map is entirely incompatible with data from different crystal forms. The `matchmaps.mr` extension of MatchMaps allows for model-bias-free comparisons of electron densities regardless of crystal form, opening up a new world of structural comparisons. For instance, an isomorphous difference map cannot characterize the impacts of crystal packing. As shown above, MatchMaps can create such a map and thus allows enhanced understanding of the often subtle role of crystal packing on protein structure.

MatchMaps depends only on the common CCP4 and Phenix crystallographic suites along with various automatically installed pure-python dependencies. MatchMaps runs in minutes on a modern laptop computer. The only required input files are a PDB or mmCIF file containing the protein model, two MTZ files containing structure factor amplitudes and uncertainties, and any CIF ligand restraint files necessary for refinement. These are the same inputs required for many common purposes (such as running `phenix.refine`) and would likely already be on hand. As outputs, MatchMaps produces real-space maps in the common MAP/CCP4/MRC format which can be readily opened in molecular visualization software such as PyMOL or Coot. For these reasons, MatchMaps should slot naturally into the crystallographer’s workflow for analysis of related datasets. Additionally, MatchMaps is open-source and can be easily modified for a new use case by an interested developer. The authors welcome issues and pull requests on GitHub for the continued improvement of the software.

## 6 Acknowledgements

We thank Harrison Wang (Harvard University) for testing of the MatchMaps code. We thank Prof. James Fraser (UCSF) and his lab for testing MatchMaps, suggesting improvements, and commenting on the pre-print of this manuscript. We thank Marcin Wojdyr (Global Phasing Ltd.) for assistance with the gemmi library. This work was supported by the NIH Director’s New Innovator Award (DP2-GM141000, to D.R.H.).

## Notes

https://doi.org/10.5281/zenodo.10452581

### Competing Interest Statement

The authors have declared no competing interest.

### Summary of Updates

Minor changes and reorganizations have been made throughout to increase clarity. Figure 1 was updated to include an additional plot where the correlations between isomorphous and poorly isomorphous data are compared per resolution bin. The manuscript now compares MatchMaps to other existing methods.

